# Endocannabinoid-dopamine interactions mediate incidental associations in the hippocampus

**DOI:** 10.64898/2026.08.24.746680

**Authors:** U.B. Fundazuri, M. Barrera-Conde, E. Rampini, P. Gómez-Sotres, C. Ioannidou, J. Pinho, M. González-Portilla, S. Beriain, A. Busquets-Garcia, G. Ferreira, G. Marsicano

## Abstract

Reinforced conditioning allows individuals predicting future events with high confidence. However, many daily behaviours rely on unreinforced connections of neutral stimuli, called Incidental Associations (IAs), which enhance predictive capacity in unstable environments and are observed across species. IAs can be studied through sensory preconditioning paradigms, where two neutral stimuli (S1/S2) are presented together in a preconditioning phase, followed by classical conditioning of S1 with a potent reinforcer. As a result, subjects present a direct response to the S1 stimulus, but also display mediated responses to the S2 stimulus never explicitly reinforced, indicating IA formation during preconditioning. Our previous work demonstrated that type-1 cannabinoid receptors (CB1 receptors) in the hippocampus are essential for this IA formation. As dopamine signaling is also important for this process, we investigate the role of interactions between these dopamine-cannabinoid systems in IA memory formation in the hippocampus.

Extending our previous work on odor-taste association, using light-sound association we showed that global CB1 receptor knock-out or specific hippocampal CB1 receptor deletion also blocked mediated responses to sound (S2) while direct response to light (S1) was unaltered. Focusing on dopamine, we then found hippocampal dopaminergic activity is enhanced during paired presentations of S1 and S2 and that blockade of dopamine D1 receptor during preconditioning S1-S2 associations abolished mediated response to S2. Interestingly, mice lacking CB1 receptors specifically in D1-receptor-expressing cells (D1-CB1-KO) failed to show mediated responses for either light-sound or odor-taste associations, identifying this CB1 receptor population as critical for IA formation. Enhanced activation of CB1 receptors, through either increase of endocannabinoids (using degradation enzyme inhibition) or exogenous stimulation by cannabis-derived Δ9-tetrahydrocannabinol (THC), was able to promote the formation of IAs under insufficient conditions. The effect of endogenous CB1 activation, but not THC, was blocked in D1-CB1-KO mice indicating that IA-facilitation by endogenous and exogenous receptor activation rely on different mechanisms. Overall, these data uncover new mechanisms underlying unreinforced associative learning.

## INTRODUCTION

Everyday decisions are guided by diverse stimuli, whose salience can ultimately determine behavior. While highly salient stimuli can directly be considered as positive or negative reinforcers, neutral stimuli are not eliciting behavioral responses per se (Treviño, 2016). However, neutral stimuli can be associated with reinforcers, acquiring new meanings that subsequently trigger behavioral responses. These kind of reinforced processes facilitate individuals adaptation to the environment (Johansen et al., 2011; Pavlov, 1927). Daily choices are often shaped by other kinds of associations, the so-called unreinforced or incidental associations (IAs), in which neutral stimuli are linked in absence of any reinforcer. Thus, IAs arise from relationships between neutral stimuli, providing non-reinforced associations that help predict outcomes (Bornstein et al., 2017; Wimmer et al., 2012; Wimmer & Shohamy, 2012). IAs extend learning beyond reinforced associations, enhancing adaptability and survival in ambiguous contexts (Busquets-Garcia & Holmes, 2022; Gewirtz & Davis, 2000; Ioannidou et al., 2021; Parkes & Westbrook, 2011). While the mechanisms of reinforcement-driven learning have been extensively characterized since Pavlov’s pioneering work (Pavlov, 1927), the processes governing IA formation remain poorly understood, despite growing evidence of their fundamental importance for human cognition and adaptive behavior (Brogden, 1947; Yu et al., 2014) and IA distortion has been implicated in psychiatric conditions such as psychosis (Busquets-Garcia, Soria-Gómez, Ferreira, et al., 2017; Busquets-Garcia, Soria-Gómez, Redon, et al., 2017; Fry et al., 2020; M. A. McDannald et al., 2011; M. McDannald & Schoenbaum, 2009; Schmack et al., 2015).

IA formation is well conserved across species, as witnessed through the use of sensory preconditioning procedures (Brogden, 1939; Ducourneau et al., 2025; Holmes & Westbrook, 2017; Kojima et al., 1998; Reid, 1952; Wong et al., 2019). In these paradigms, two neutral stimuli (S1/S2) are presented together during a preconditioning phase, followed by classical conditioning of S1 with a potent reinforcer. As a result, subjects display a direct response to S1, but also show mediated responses to S2, despite never having been directly reinforced, revealing the formation of IA during preconditioning (Brogden, 1939; Gewirtz & Davis, 2000; Holmes et al., 2022; Ward-Robinson et al., 2005). Among the different brain regions involved in sensory preconditioning (González-Parra et al., 2025; Holmes et al., 2013; Sadacca et al., 2018; Ward-Robinson et al., 2005; Wong et al., 2019, 2026), the hippocampus has been historically proposed to play a key role across conditioning and retrieval phases (Iordanova et al., 2011; Wheeler et al., 2013; Wimmer & Shohamy, 2012; Talaron et al., 2026). However, recent studies have also demonstrated that this region is relevant for IA formation during the preconditioning phase itself, when the two neutral stimuli are first paired (Busquets-Garcia et al., 2018; Pinho et al., 2025; Talaron et al., 2026). Notably, we have previously shown that the hippocampal endocannabinoid system (ECS), through its main type-1 cannabinoid receptors (CB1 receptors), is required for IA formation (Busquets-Garcia et al., 2018). Deletion of hippocampal CB1 receptors or their blockade specifically during S1-S2 association both abolished mediated responses to S2, leaving direct response intact. Another neuromodulatory system, the dopamine (DA) system, plays a critical role in sensory preconditioning (Roughley et al., 2021; Sharpe et al., 2017), and hippocampal DA signaling has been implicated in memory linking, a complex memory process that shares features with IAs (Chowdhury et al., 2022; Seitz et al., 2021). Interestingly, the ECS is well known to regulate other neurotransmitter systems including the DA system (El Khoury et al., 2012; García et al., 2016) and interactions between DA and ECS have been shown to regulate a variety of complex behavioral and cognitive processes (Diana et al., 1998; Laricchiuta et al., 2014; Luján et al., 2023, 2026; Melis et al., 2004; Melis & Pistis, 2007; Terzian et al., 2011). Yet, the potential role of hippocampal DA-ECS interactions in IA formation remain unknown.

Building on our previous research on odor-taste association (Busquets-Garcia et al., 2018), we first investigated whether hippocampal CB1 receptors govern IA formation using light-sound association. We then investigated the potential interaction between the endocannabinoid and dopaminergic systems during IA formation combining fiber photometry, pharmacological and genetic approaches, showing that both systems are key players in IA memory formation.

## MATERIALS AND METHODS

All experiments were conducted in strict compliance with the European Union recommendations (2010/63/EU) and were approved by the French Ministry of Agriculture and Fisheries (authorization number B33063098) and the local ethical committee (authorization APAFIS#44909).

### Animals

C57BL/6-N (Janvier, France) and inbred constitutive CB1 mutant (center’s facility, with a C57BL/6-N background), both male and female mice (*Mus musculus*) were used in this study. CB1 mutant lines included: CB1^f/f^ mice (CB1-flox), carrying a floxed version of the CB1 gene and also used as wild-type (WT) controls (Marsicano et al., 2003); CB1 receptor knockout mouse line (CB1-KO) carrying a constitutive global deletion of the CB1 gene (Marsicano et al., 2002); and the D1-CB1 knockout mouse line (D1-CB1-KO) carrying a conditional deletion of the CB1 gene in Dopamine D1 receptor-positive cells under the control of Drd1-Cre recombinase (Monory et al., 2007). WT littermates of each line were respectively used as controls for the behavioral experiments. Offspring heterozygous for transgenes were genotyped following established protocols. All animals are housed in groups of two to four, with *ad libitum* access to food and standard enrichment materials, with a 12-hour light/dark cycle (lights on at 7am and off at 7pm). Experiments were conducted in the light phase from 9am to 5pm.

### Light-sound preconditioning protocol

The light and sound sensory preconditioning protocol was adapted from the study of Pinho et al. (2025) due to different mouse housing conditions (light vs dark phases inverted). We therefore adjusted light and sound intensities and context to the conditions in our animal facility. The task was performed in automated conditioning poly-boxes (Imetronic, France), each measuring 30 × 40 × 36 cm and consisting in a light- and sound-attenuated cabinet with a floor made of metal rods. One of the walls is transparent, allowing video recording of every phase of the protocol.

#### Habituation

On the first day of the protocol (day 1), animals were placed in the chamber for a single 20-minute session.

#### Preconditioning (Phase 1)

This phase consisted of 6 sessions over 3 consecutive days (days 2-4, 2 sessions per day with a minimum intersession interval of 3 hours). For the paired groups, every session began with a 3-minutes OFF period followed by five simultaneous 30-second exposures to a light (S1, house light, located on top of the chamber) and a sound (S2, 50 dB, 1000 Hz), with a 30-second intertrial interval (ITI), and ended with a 1-minutes OFF period, for a total session duration of 8 minutes and 30 seconds. This resulted in a total of 30 simultaneous light-sound presentations during this phase.

For the unpaired group, light and sound were never presented together within the same session, but rather in separate preconditioning sessions in a pseudorandomized order: S1 light, S2 sound, S3 sound, S4 light, S5 light and S6 sound. These sessions had the same duration as the paired sessions, and animals were exposed to each stimulus for the same total amount of time.

For the reduced preconditioning group (rPC), mice underwent only 2 sessions within the same day (day 2, with a minimum intersession interval of 3 hours). Sessions were identical to those described before, except that mice received a total of 10 simultaneous light-sound presentations. The following phases happen as afterwards described in Phase 2 and Phase 3 in days 3 and 4 respectively.

#### Conditioning (Phase 2)

Conditioning consisted of two sessions on the same day (day 5, minimum intersession interval of 3 hours). Paired and unpaired groups underwent identical conditioning sessions lasting 8 minutes and 50 seconds of duration. Each session began with a 3-minutes OFF period, followed by 5 light exposures of 10 seconds co-terminating with a mild footshock (0.4 mA, 2 seconds), with a 1-minute ITI, and ending with a 1-minute OFF period.

#### Tests (Phase 3)

The sound test (mediated responses test) was performed during the morning (day 6), followed by the light test (direct responses test, minimum intersession interval of 1 hour). The context in this phase differed from the previous ones: a new chamber, with different wall colors and floor texture, was introduced in the poly-boxes, and an odor was added while cleaning the chamber (Phagospray, Christeyns, France). Both tests lasted 6 minutes and consisted of a 3-minute OFF period followed by a 3-minute ON period, during which the sound (mediated response test) or the light (direct response test) was continuously presented.

Freezing (i.e., immobility) was used as the behavioral readout. Video recordings enabled automatic quantification of freezing using a custom Python script, available at: https://github.com/paulagsotres/still_count. Freezing was defined as a change of lower than 50 pixels between frames, with a minimum of 4 consecutive frames required for a bout to be counted as freezing (videos were recorded at 24.67 fps). Automatic counts were validated against manual counts performed by blinded expert observers, and a correlation of more than 0.95 was observed.

To calculate the behavioral responses, we considered the last minute of the OFF period (seconds 120-180) and the first minute of the ON period (seconds 180-240) of each test. The presence of mediated and direct responses was determined by comparing the freezing percentage during the ON versus OFF phase, and by computing a freezing index as: (Freezing ON – Freezing OFF)/(Freezing ON + Freezing OFF). Mice in control and WT groups with a freezing percentage >40% in the OFF period of the mediated responses test were excluded from analysis.

For pharmacological experiments with SCH23390 (0.03 mg/kg), mice were habituated to injections for 2 days during the week preceding the behavioral experiment. The drug and its vehicle were administered i.p. 15 minutes before each preconditioning session.

For pharmacological administration of JZL195 (8 mg/kg) and Δ9-tetrahydrocannabinol (THC, 1 mg/kg), mice were likewise habituated to injections for 2 days during the preceding week. Both drugs and their respective vehicles were injected i.p. 2 hours before each preconditioning session in Phase 1.

### Odor-taste preconditioning protocol

Mice were group-housed in the same room where the protocol took place. Each day of the protocol, animals were individually placed in another housing cage and they had 1-hour access to a bottle containing the odor/taste mix, as previously described (Busquets-Garcia et al., 2018; Busquets-Garcia et al., 2017a, 2017b). The solutions used in this sensory preconditioning task were presented in 50-mL drinking bottles with either banana (0.05%, isoamyl acetate) or almond (0.01%, benzaldehyde) as odors, and sucrose (5%) or maltodextrin (5%) as tastes. All compounds were obtained from Sigma-Aldrich (St. Quentin Fallavier Cedex, France).

#### Habituation

Water deprivation began 24 hours before the start of the protocol. All subjects then received 1-hour access to water for 3 consecutive days as habituation. Mice that drank <1 mL were given an extra hour to ensure habituation to the drinking procedure.

#### Preconditioning (Phase 1)

This stage consisted of 3 odor-taste pairings across 6 days. Each pairing corresponded to two days: on the first day, subjects had 1-hour access to a flavored solution containing a new taste (5% sucrose) and a new odorant (0.05% banana) mixed with water. On the second day, animals received a taste (5% maltodextrin) and an odor not given the previous day (0.01% almond).

#### Conditioning (Phase 2)

Conditioning consisted of 6 days, during which the banana odor was devalued to become the conditioned stimulus. On days 1, 3, and 5 of this phase, subjects received 1-hour access to almond. On days 2, 4, and 6, mice received 1-hour access to banana, immediately followed by an i.p. injection of lithium chloride (LiCl, 0.3 M, 1% body weight), which induces gastric malaise in mice. After conditioning, subjects underwent a recovery day during which they had 1-hour access to water.

#### Tests (Phase 3)

Over the next 2 days, mediated and direct responses were assessed using a 1-hour two-choice test. The mediated response was evaluated on the first test day with a two-choice test between the sucrose taste (previously associated with the banana odor but never directly paired with LiCl) and the maltodextrin taste (previously associated with almond). On the second test day, the direct response was measured with a two-choice test between the two odors.

For this sensory preconditioning paradigm, data were presented as absolute liquid intake for each odor or taste. The presence of mediated responses was determined by comparing maltodextrin intake to sucrose intake, or by the consumption index: (Maltodextrin – Sucrose)/(Maltodextrin + Sucrose). Direct responses were evaluated by comparing almond odor consumption to banana odor intake, or by calculating the consumption index: (Almond – Banana)/(Almond + Banana).

### Drug preparation and administration

SCH23390 (R(+)-SCH-23390 hydrochloride, ≥98%) was purchased from Sigma-Aldrich (St Quentin Fallavier, France) and dissolved in saline (NaCl 0.9%, Fisher Scientific, Illkirch Cedex, France) at a dose of 0.03 mg/kg. JZL195 (8 mg/kg) was purchased from Tocris Bioscience (Bristol, United Kingdom) and dissolved in 2.5% DMSO (dimethyl sulfoxide, sterile-filtered, Sigma-Aldrich, St Quentin Fallavier, France), 2.5% Kolliphor-EL (Sigma-Aldrich, St Quentin Fallavier, France) and 95% saline (NaCl 0.9%). Δ^9^- tetrahidrocannabinol (THC, 1 mg/kg) was purchased from HC-Pharm (Frankfurt, Germany) and dissolved in 5% absolute ethanol, 4% Kolliphor-EL and 91% saline (NaCl 0.9%).

### Surgery for viral injection and fiber implantation

Mice were injected intraperitoneally (i.p.) with buprenorphine (0.05 mg/kg, Buprecare), anesthetized using 5% isoflurane, and placed into a stereotaxic apparatus (Model 900, Kopf Instruments, CA, USA; with mouse adaptor and lateral ear bars), maintained under 2% isoflurane for the duration of the surgery. Local analgesia with lidocaine (0.1 ml at 0.5%, Lidor) was applied under the skin of the head before incision.

To assess the specific contribution of neuronal CB1 receptors in the hippocampus to light and sound sensory preconditioning, CB1-flox mice were injected with viruses purchased from ETH (Zurich, Switzerland): AAV-hSyn-CRE-GFP (v146-8) or its control AAV-hSyn-mCherry (v133-8), expressing the mCherry reporter in neurons. Viral injections for CB1 receptor deletion were delivered bilaterally in the hippocampus through a microsyringe (0.25 mL Hamilton syringe with a 30-gauge beveled needle) attached to a pump (UMP3-1, World Precision Instruments, FL, USA). In all surgeries, mice received two injections per site (0.5 μl each) bilaterally, at the following coordinates according to Paxinos and Franklin (Paxinos & Franklin, 2001): AP – 2; ML ± 1.5; DV – 2 (1^st^ injection) and – 1.5 (2^nd^ injection) at a speed of 250 nl/min.

To study in vivo dopamine signaling in hippocampal neurons by fiber photometry during light and sound sensory preconditioning, the same surgical procedure described above was followed. However, mice were injected unilaterally with the AAV-hSyn-GRAB(DA2m) viral construct (Sun et al., 2020), kindly provided by Pierre Trifilieff and Lola Hardt (Nutrineuro, Bordeaux, France). We used a total volume of 0.8 μl, at the following coordinates: AP –2; ML –1.5; DV –1.7, at a speed of 250 nl/min. The optical fiber (400 μm diameter, 0.5 NA, 2 mm long, RWD, China) was then placed 200 μm above the injection site (i.e., at DV –1.5) and fixed with dental cement (Super-Bond Universal Kit, SUN MEDICAL, Japan). All viruses were titered between 2–8 × 10¹¹ genomic copies/mL.

Following surgery, all mice received an i.p. injection of 0.2 ml saline solution and the anti-inflammatory drug meloxicam (5 mg/kg, Metacam), continued for 2 additional days. Animals continued to be housed collectively, and body weight was monitored daily for 4–5 days to assess recovery. Behavioral and fiber photometry experiments were both carried out 4–5 weeks after surgery.

### Fiber photometry experiments

Mice with fiber implants were habituated to being connected to the fiber photometry cable for 2 minutes per day, over 3 days prior to the experiment. For the dopamine GRAB(DA2m) sensor, the fiber photometry set-up collected emitted fluorescence with an sCMOS camera (Hamamatsu Orca Flash v3) through an optic fiber (core 400 μm, N.A. 0.5) divided into 2 sections: a short fiber implanted in the mouse’s brain and a long fiber (modified patch cord), connected via a ferrule-ferrule (1.25 mm) connection. To minimize photobleaching during recording and preserve a high signal-to-noise ratio, light intensity at the tip of the patch cord was adjusted to ∼100 μW for the 470 nm channel and ∼30 μW for the 405 nm (isosbestic) channel. A custom MATLAB script (MathWorks) was used to synchronize video recording with fiber photometry, together with a programmed Arduino board. The sampling rate was set to 20 Hz for both photometry (interleaved) and video recording.

Dopamine signals were recorded during Phases 1–3 of the light and sound sensory preconditioning paradigm, with mice exposed to either paired or unpaired preconditioning (Phase 1). The signal was pre-processed and processed using a custom Python script, available at: https://github.com/paulagsotres/Gomez-Sotres-FP-analysis-GUI. Raw dopamine signals were pre-processed by removing the first 2 minutes of the recording, to reduce the effect of the initial exponential photobleaching, and by removing point artifacts. The signal was filtered using a high-pass filter (0.01 Hz) and a low-pass filter (3 Hz). The 470 nm signal was fitted to the isosbestic 405 nm signal using airPLS (polynomial order 4). The normalized fluorescence change (ΔF/F) was calculated at each time point as (F470−F405,fitted)/F405,fitted. For each of the events per recording, the ΔF/F trace was extracted together with a 10 s pre-event baseline, and this extracted window (baseline + response) was z-scored using statistics computed from the window itself.

For preconditioning (Phase 1), the signal corresponding to the 35 s following cue onset was extracted, with a 10 s baseline preceding onset, and z-scored as described above. The first positive peak in following cue onset in the z-scored data was used to compare responses to the cues between paired (simultaneous light-sound exposure) and unpaired (separate light and sound exposure) groups.

In conditioning (Phase 2), the signal corresponding to the 25 s following light onset was extracted, with a 10 s baseline preceding onset, and z-scored as described above. The first positive peak following shock onset in the z-scored data was used to compare responses to the shock between groups.

For both the mediated response test (sound) and the direct response test (light), the signal corresponding to the 185 s following the onset of each cue was extracted, with a 10 s baseline preceding cue onset, and z-scored as described above. The first positive peak following cue onset in the z-scored data was used to compare responses to the cues between groups.

### Immunostaining for light microscopy

AAV-injected mice were deeply anesthetized with an intraperitoneal injection of xylazine (20 mg/kg), followed by euthasol (400 mg/kg). Mice were transcardially perfused with 20 mL of phosphate-buffered saline (PBS, 0.1 M, pH 7.4) for 2 min, followed by 50 mL of 10% wt/vol neutral buffered formalin (Sigma, HT501128-4L) for 5 min. After perfusion, brains were extracted and post-fixed in 10% wt/vol neutral buffered formalin for 24 h. Brains were then transferred to a 30% wt/vol PBS-sucrose solution for cryopreservation. Once brains were fully dehydrated and had sunk to the bottom of the tube (on average, 3–5 days), they were frozen in isopentane and cut into 30 μm coronal sections using a cryostat (Leica Biosystems, CM1950S). Hippocampal slices were stored in antifreeze solution at −20 °C until further use.

All sections were stained with 4′,6-diamidino-2-phenylindole (DAPI, 1:20000, Invitrogen D3571) to visualize cell nuclei, washed with PBST (0.3% Triton X-100 in 1X PBS, pH 7.4), and then mounted, dried, and coverslipped. Sections were examined using a SlidesVIEW VS200 (Evident Scientific) scanner to verify intrinsic fluorescence of the viruses and correct placement of the optic fiber. Mice whose brains did not meet expression requirements, or that showed optic fiber misplacement, were excluded from the experiments. Micrographs were acquired as a maximum projection of 7 stack 40x images and we created a composite to get the whole hippocampus.

### Data collection and statistical analysis

Statistical methods were not used to predetermine sample size; however, the number of animals used was comparable to that reported in previous studies (Busquets-Garcia et al., 2017; Busquets-Garcia et al., 2018; Pinho et al., 2025). Mice were randomly assigned to the experimental conditions. For the fiber photometry experiments, signals were extracted and processed as described above using the provided custom code. Freezing behavior in the light-sound sensory preconditioning paradigm was analyzed using custom software. Raw data from all experiments were processed and analyzed using Microsoft Excel 2021 and GraphPad Prism 11.0.2.

Graphs and statistical analyses were generated using GraphPad Prism 11.0.2. All data represent independent biological replicates (individual mice) and are presented as individual data points with the mean ± standard error of the mean (SEM). Data normality was assessed using the Shapiro–Wilk test. Depending on the outcome, either parametric tests (Student’s t-test, one-sample t-test, ordinary one-way ANOVA followed by Tukey’s multiple-comparisons test, repeated-measures two-way ANOVA followed by Sidak’s multiple-comparisons test, two-way ANOVA followed by Tukey’s multiple-comparisons test, or three-way ANOVA followed by Tukey’s multiple-comparisons test) or non-parametric tests (Mann–Whitney test or one-sample Wilcoxon signed-rank test) were performed.

## RESULTS

### Hippocampal CB1 receptors are required for light-sound sensory preconditioning

To evaluate whether CB1 receptors control light-sound-based IAs as previously shown for odor-taste association (Busquets-Garcia, Oliveira Da Cruz, et al., 2018), we adapted in both male and female mice a recent protocol described by Pinho et al. (2025) (Fig. 1A, see Methods). Animals were assigned to either a paired group, in which light and sound cues were presented simultaneously during the preconditioning phase, or an unpaired control group, in which both cues were presented separately. Both groups received a subsequent light fear conditioning and were tested for freezing response to light (direct response) and sound (mediated response) quantified automatically using a custom Python-based algorithm.

**Figure 1:**
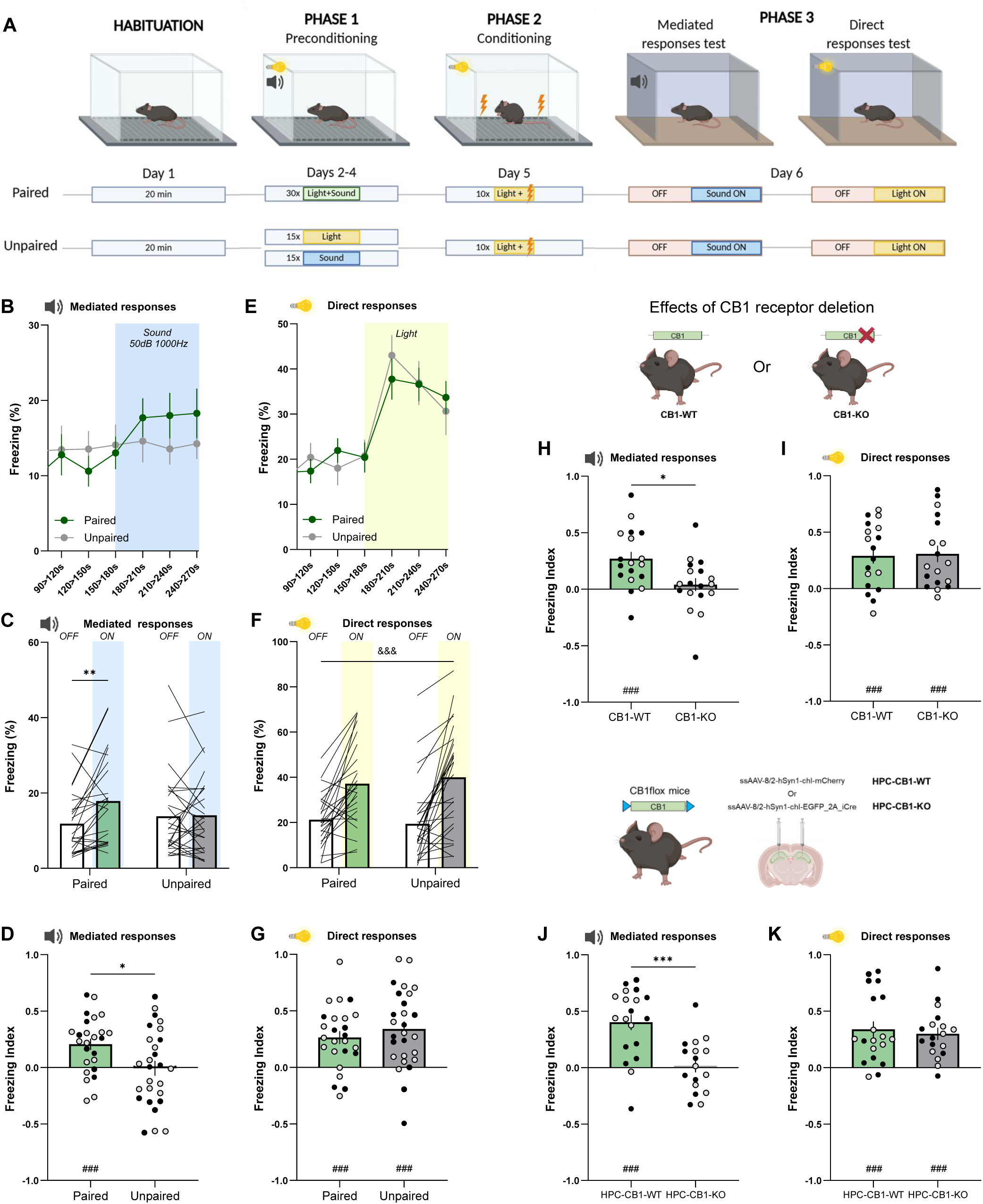
Hippocampal CB1 receptors are required for light sound sensory preconditioning. (A) Schematic representation of the light and sound sensory preconditioning task with the paired and unpaired preconditioning experimental groups. (B-G) (B) Temporal line of the freezing percentage -in bins of 30 seconds- of the paired and unpaired experimental groups in the 90 seconds before and after 90 seconds of sound display in the mediated responses test of the paradigm. (C) Freezing percentage of the paired and unpaired groups in the last 60 seconds of the OFF period and the first 60 seconds of the ON period in the sound test. Repeated measures two-way ANOVA, interaction p=0,0230; Sidak’s post-hoc, p(ON vs OFF in the paired group)=0,0026. (D) Freezing index of the paired and unpaired groups in the mediated responses test. Student’s unpaired t-test, p=0.0133. One sample t-test, p(paired)=0.0003. (E) Temporal line of the freezing percentage -in bins of 30 seconds-of the paired and unpaired experimental groups in the 90 seconds before and after 90 seconds of sound display in the direct responses test of the paradigm. (F) Freezing percentage of the paired and unpaired groups in the last 60 seconds of the OFF period and the first 60 seconds of the ON period in the direct responses test. Repeated measures two-way ANOVA, ON vs OFF effect p<0.0001. (G) Freezing index of the paired and unpaired groups in the direct responses test. One sample t-test, p(paired)<0.0001, p(unpaired)<0.0001. (H and I) Freezing index after the paired preconditioning in CB1-WT and CB1-KO mice in the mediated responses (H) and direct responses (I) tests. (H) Student’s unpaired t-test, p=0.0103. One sample t-test, p(CB1-WT)=0.0004. (I) One sample t-test, p(CB1-WT)=0.0006, p(CB1-KO)=0,0006. n_(CB1-WT)_= 10M/8F, n_(CB1-KO)_= 9M/9F. (J and K). Freezing index after the paired preconditioning in HPC-CB1-WT and mice with hippocampal CB1 receptor deletion (HPC-CB1-KO) in the mediated responses (J) and direct responses (K) tests. (J) Student’s unpaired t-test, p=0.0002. One sample t-test, p(HPC-CB1-WT)<0.0001. (K) One sample Wilcoxon test, p(HPC-CB1-WT)<0.0001. One sample t-test p(HPC-CB1-KO)<0.0001. n_(HPC-CB1-WT)_= 11M/8F, n_(HPC-CB1-KO)_= 8M/9F. Black dots: males; grey dots: females. Data are expressed in mean ± SEM. *p<0.05, **p<0.01, ***p 0.001, ^###^p<0.001, ^&&&^p<0.001.

After undergoing light/sound paired or unpaired preconditioning and light fear conditioning, sound presentation induced a freezing increase only in the paired group (e.g. mediated responses), with significantly higher freezing during the first minute of the ON period compared with the last minute of the OFF period (Fig. 1B,C). The freezing index was also higher in the paired group relative to the unpaired group (Fig. 1D), with no differences between male and female mice in either group (Supplementary Fig. 1A). When light was presented, both experimental groups showed similar direct responses, reflected in enhanced freezing from the OFF to the ON period (Fig. 1E,F). The freezing index was similar in both groups (Fig. 1G) or sexes (Supplementary Fig. 1B). Taken together, these results indicate that only simultaneous, paired presentations during preconditioning induce mediated responses in both male and female mice.

We then evaluated whether CB1 receptors control light-sound-based IAs as previously shown for odor-taste association (Busquets-Garcia et al., 2018). Using CB1-KO male and female mice (Marsicano et al., 2002), we found impaired mediated responses (Fig. 1H), but normal direct responses (Fig. 1I). These results demonstrate the necessity of CB1 receptors in light-sound preconditioning, consistent with previous findings in the odor-taste paradigm (Busquets-Garcia et al., 2018). As the hippocampus has been identified as a key region where CB1 receptors play a pivotal role in odor-taste preconditioning (Busquets-Garcia et al., 2018), we then determine whether hippocampal CB1 receptors are involved in light-sound preconditioning. For this, an AAV carrying Cre-recombinase (AAV-hSyn-CRE-GFP) was infused into both hippocampal subregions of CB1-flox male and female mice, to achieve region-specific CB1 receptor deletion (HPC-CB1-KO). This manipulation did not alter direct responses to light (Fig. 1K) but HPC-CB1-KO mice showed impaired mediated responses to sound. No differences were found between sexes (Supplementary Fig. 1H,J). These results demonstrate that hippocampal CB1 receptors are required for mediated responses regardless of sensory modalities used during preconditioning.

### Hippocampal dopamine signaling is necessary to form IAs

The ECS is not the only neuromodulatory system controlling IAs. The dopaminergic system has been shown to play a role in IAs and related processes (Costa et al., 2025; Roughley et al., 2021; Sharpe et al., 2017, 2020; Young et al., 1998), identifying the hippocampus as an important dopaminoceptive region for memory linking (Chowdhury et al., 2022). To characterize hippocampal dopamine activity, we injected in the hippocampus of wild-type mice an AAV coding for the GRAB(DA2m) sensor (Sun et al., 2020) and implanted an optic fiber. We then recorded hippocampal dopamine activity in freely moving mice using *in vivo* fiber photometry during light-sound preconditioning, light conditioning and mediated and direct responses (Fig. 2A). Animals were divided into two experimental groups: a paired group, receiving simultaneous light and sound presentations during preconditioning, and an unpaired group, which received separate light and sound exposure (see Methods). Hippocampal dopamine activity was then quantified as z-scored ΔF/F, spanning the 10 s before and after stimuli onset, considering the peak z-scored ΔF/F following light, sound or footshock onset.

**Figure 2:**
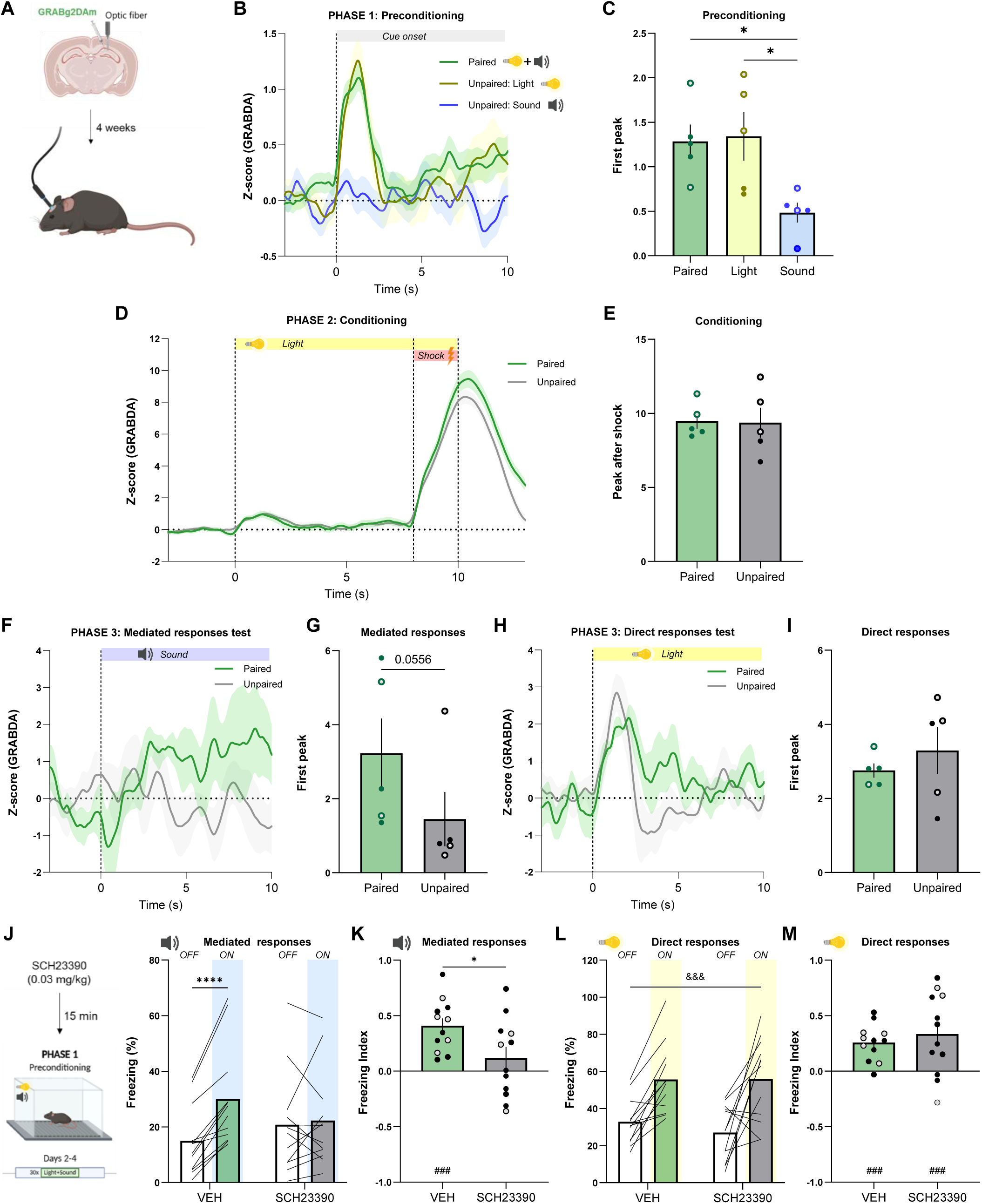
Hippocampal dopamine signaling is necessary to form IAs. (A) Schematic representation of Dopamine sensor injection and optic fiber implant for later recordings in the mouse hippocampus. (B-I) (B) Average Z-scored ΔF/F Dopamine responses in neurons of the hippocampus aligned to the 30 light and sound simultaneous presentation onset in mice following paired preconditioning in the Phase 1 of the paradigm, or aligned to the 15 sound and 15 light presentations in mice following unpaired preconditioning. (C) First peak of average Z-scored ΔF/F Dopamine responses In the paired and unpaired (light and sound separated) groups. Ordinary one-way ANOVA, p=0.019; Tukey’s post-hoc, p(Paired vs Sound)=0.04, p=(Light vs Sound)=0.0278. (D) Average Z-scored ΔF/F Dopamine responses aligned to the 10 light and shock presentation onset during conditioning for the paired and unpaired groups. (E) First peak of average Z-scored ΔF/F Dopamine responses in the paired and unpaired groups after the footshock. (F) Z-scored ΔF/F Dopamine responses aligned to the sound presentation onset during the mediated responses test in the Phase 3 of the protocol for the paired and unpaired groups. (G) First peak of Z-scored ΔF/F Dopamine responses in the paired and unpaired groups after the sound onset. Mann Withney test, p=0.0556. (H) Z-scored ΔF/F Dopamine responses aligned to the light presentation onset during the direct responses test for the paired and unpaired groups. (I) First peak of Z-scored ΔF/F Dopamine responses in the paired and unpaired groups after the light onset. n_(paired)_= 3M/2F, n_(unpaired)_= 2M/3F. (J-M) (J) Schematic representation of SCH23390 (0.003 mg/kg) injections 15 minutes before each preconditioning session. Freezing percentage of the SCH23390 (0.03 mg/kg) and vehicle-treated mice in the last 60 seconds of the OFF period and the first 60 seconds of the ON period in the sound test. Repeated measures two-way ANOVA, interaction p=0.0038; Sidak’s post-hoc, p(ON vs OFF in the vehicle group)<0.0001. (K) Freezing index after the paired preconditioning of SCH23390 (0.03 mg/kg) and vehicle-treated mice in the mediated responses test. Student’s unpaired t-test, p=0.0255. One sample t-test, p(vehicle)<0.0001. (L) Freezing percentage of the SCH23390 (0.03 mg/kg) and vehicle-treated mice in the last 60 seconds of the OFF period and the first 60 seconds of the ON period in the light test. Repeated measures two-way ANOVA, ON vs OFF effect p<0.0001. (M) Freezing index after the paired preconditioning of SCH23390 (0.03 mg/kg) and vehicle-treated mice in the mediated responses direct responses test. One sample t-test, p(vehicle)=0.0086, p(SCH2330)=0.0002. n_(vehicle)_= 7M/5F, n_(SCH23390)_= 9M/3F. Black and filled dots: males; grey dots: females. Data are expressed in mean ± SEM. *p<0.05, ****p<0.001, ^###^p<0.001, ^&&&^p<0.001.

During preconditioning, the average dopamine peak in response to the paired light-sound cues (paired group) and to the light alone (unpaired group) was significantly higher than the peak in response to the sound alone (unpaired group; Fig. 2B,C). This pattern was consistent across each preconditioning session (Supplementary Fig. 2K), resulting in differential dopamine activity across preconditioning days when comparing paired and unpaired groups (Supplementary Fig. 2L). Altogether, these data suggest that hippocampal dopamine encodes light exposure and is consistently higher in the paired group. During conditioning, we did not observe any difference in peak amplitude after footshock between paired and unpaired groups (Fig. 2D,E). However, during the mediated response to sound, the first hippocampal dopamine peak was higher in the paired group (Fig. 2F,G). This difference was not observed for the direct responses to light (Fig. 2H,I). This suggests that hippocampal dopamine activity is specifically enhanced during the mediated responses to the sound after light-sound pairing.

To evaluate whether dopamine activity during preconditioning is important for future mediated responses, we blocked dopamine receptor before each preconditioning session by administering a dopamine D1 receptor antagonist SCH23390 (0.03 mg/kg i.p.). This treatment specifically blocked mediated responses, without affecting the direct responses (Fig. 2J–M), demonstrating that dopamine signaling through D1 receptors is required for IA formation.

### CB1 receptors in D1-positive cells are required for sensory preconditioning

Given that our data showed that the CB1 receptors and D1 receptors are both required for IA formation, we asked whether these two systems interact in this process. It is well established that CB1 receptors are expressed in D1-positive cells across a broad range of cellular populations in different brain regions (Hermann et al., 2002; Martín et al., 2008; Puighermanal et al., 2017). Mice lacking CB1 receptors in D1-positive cells show deficits in various behavioral tests (Monory et al., 2007; Terzian et al., 2011), including unreinforced learning (Oliveira Da Cruz et al., 2020). We therefore tested this mouse line in both light-sound and odor-taste preconditioning protocols. In the light-sound preconditioning, mediated but not direct responses were impaired in D1-CB1-KO male and female mice (Fig. 3A–D). We then used odor-taste preconditioning protocol (Busquets-Garcia et al., 2018; Busquets-Garcia et al., 2017a, 2017b; Wheeler et al., 2013), which consists of a presenting two different odor-taste pairs during preconditioning (banana-sucrose / almond-maltodextrin), followed by devaluation of one odor (banana) using the malaise-inducing drug lithium chloride. The mediated responses were evaluated by a choice between the two tastes (sucrose versus maltodextrin) and the direct responses by a choice between the two odors (banana versus almond; Fig. 3E; see Methods for details). In D1-CB1-KO mice, the mediated responses were abolished whereas the direct responses were similar to those of D1-CB1-WT littermates (Fig. 3H– I). These results demonstrate that CB1 receptors in D1-positive cells mediates sensory preconditioning regardless of the nature of the cues paired.

**Figure 3:**
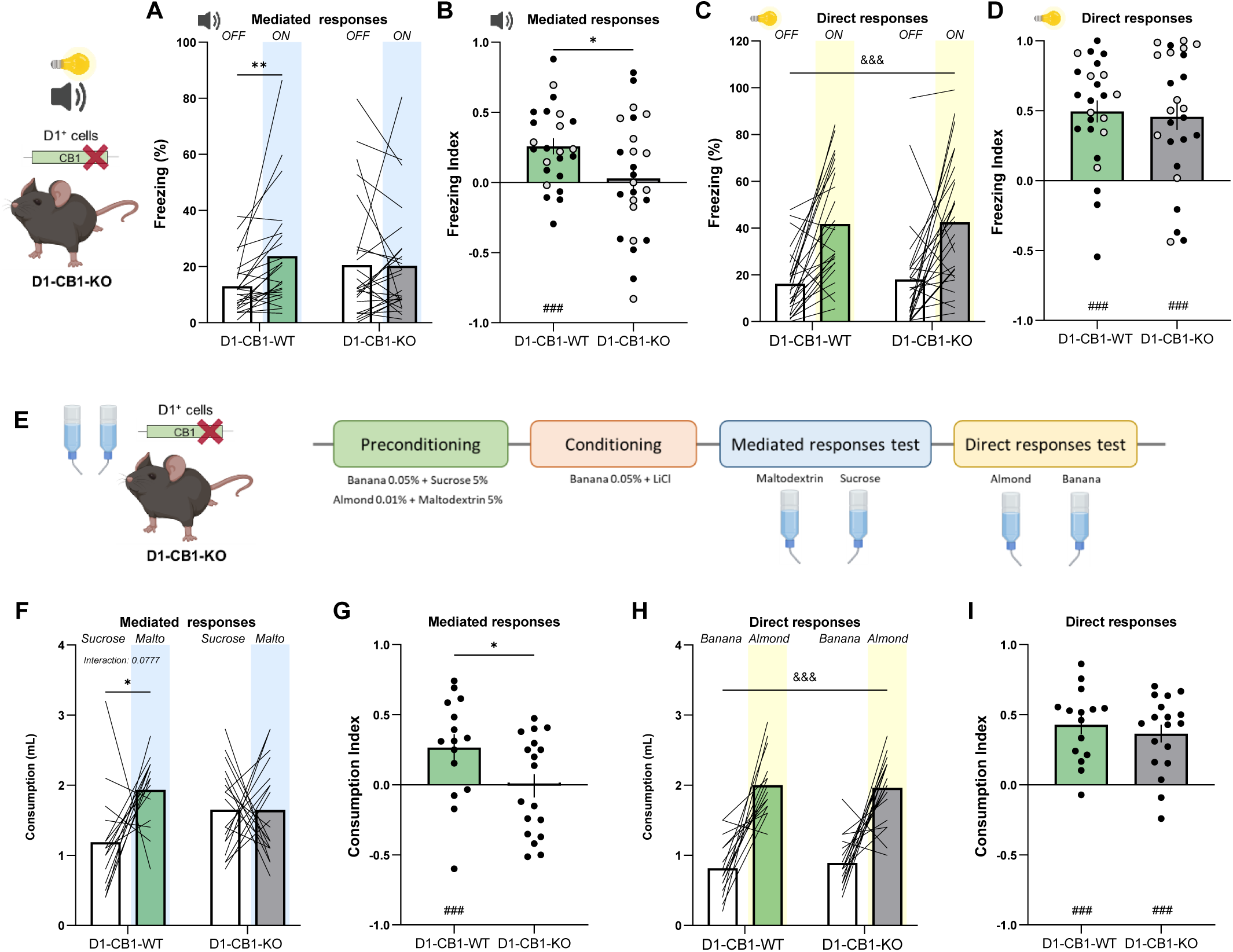
CB1 receptors in D1-positive cells are required for sensory preconditioning. (A-D) (A) Freezing percentage of the D1-CB1-WT and D1-CB1-KO groups in the last 60 seconds of the OFF period and the first 60 seconds of the ON period in the sound test. Repeated measures two-way ANOVA, interaction p=0.031; Sidak’s post-hoc, p(ON vs OFF in the D1-CB1-WT group)=0.0083. (B) Freezing index after the paired light and sound preconditioning in D1-CB1-WT and D1-CB1-KO mice in the mediated responses test. Student’s unpaired t-test, p=0.0308. One sample t-test, p(D1-CB1-WT)=0.0002. (C) Freezing percentage of the D1-CB1-WT and D1-CB1-KO groups in the last 60 seconds of the OFF period and the first 60 seconds of the ON period in the light test. Repeated measures two-way ANOVA, ON vs OFF effect p<0.0001. (D) Freezing index after the paired light and sound preconditioning in D1-CB1-WT and D1-CB1-KO mice in the direct responses test. One sample t-test, p(D1-CB1-WT)<0.0001. One sample Wilcoxon test, p(SCH2330)=0.0002. n_(D1-CB1-WT)_= 16M/8F, n_(D1-CB1-KO)_= 13M/12F. (E) Schematic representation of the light and sound sensory preconditioning task. (F-I) (F) Liquid consumption of D1-CB1-WT and D1-CB1-KO mice in the taste test in the odor and taste sensory preconditioning. Repeated measures two-way ANOVA, interaction p=0.0777; Sidak’s post-hoc, p(Sucrose vs Maltodextrin in the D1-CB1-WT group)=0.0396. (G) Consumption index after odor and taste sensory preconditioning task of the D1-CB1-WT and D1-CB1-KO mice in the mediated responses test. Student’s unpaired t-test, p=0.0363. One sample t-test, p(D1-CB1-WT)=0.0136. (H) Liquid consumption of D1-CB1-WT and D1-CB1-KO mice in the direct responses test in the odor and taste sensory preconditioning. Repeated measures two-way ANOVA, Banana vs Almond effect p<0.0001. (I) Consumption index after odor and taste sensory preconditioning task of the D1-CB1-WT and D1-CB1-KO mice in the direct responses test. One sample t-test, p(D1-CB-KO)<0.0001, p(D1-CB1-KO)<0.0001. n_(D1-CB1-WT)_= 15M, n_(D1-CB1-KO)_= 18M. Black dots: males; grey dots: females. Data are expressed in mean ± SEM. *p<0.05, **p<0.01, ^###^p<0.001, ^&&&^p<0.001.

### Endogenous and exogenous cannabinoids facilitate mediated responses through different mechanisms

It has been demonstrated that a minimal number of S1-S2 pairings during preconditioning is required to trigger mediated responses (Busquets-Garcia et al., 2017; Hoffeld et al., 1960). Therefore, we wondered whether increasing CB1 receptor activity during an insufficient number of SI-S2 pairings could promote mediated responses. Using 10 light-sound pairings (instead of the usual 30 pairings; Fig. 4A; see Methods for details), we observed that animals did not show mediated responses (Fig. 4B–E, VEH group). In order to increase CB1 receptor activity we injected WT male and female mice prior to this 10 light-sound preconditioning either JZL195 (8 mg/kg), which blocks the enzymes responsible for endocannabinoid degradation leading to increasing endocannabinoid levels and CB1 receptor activity (Long et al., 2009), or Δ9-tetrahydrocannabinol (THC, 1 mg/kg), which exogenously increases CB1 receptor activity (Devane et al., 1988); Fig. 4A; see Methods for details). Both JZL195 and THC promoted mediated responses to sound without affecting direct responses to light (Fig. 4B–E), indicating that CB1 receptor activation can facilitate IA formation.

**Figure 4:**
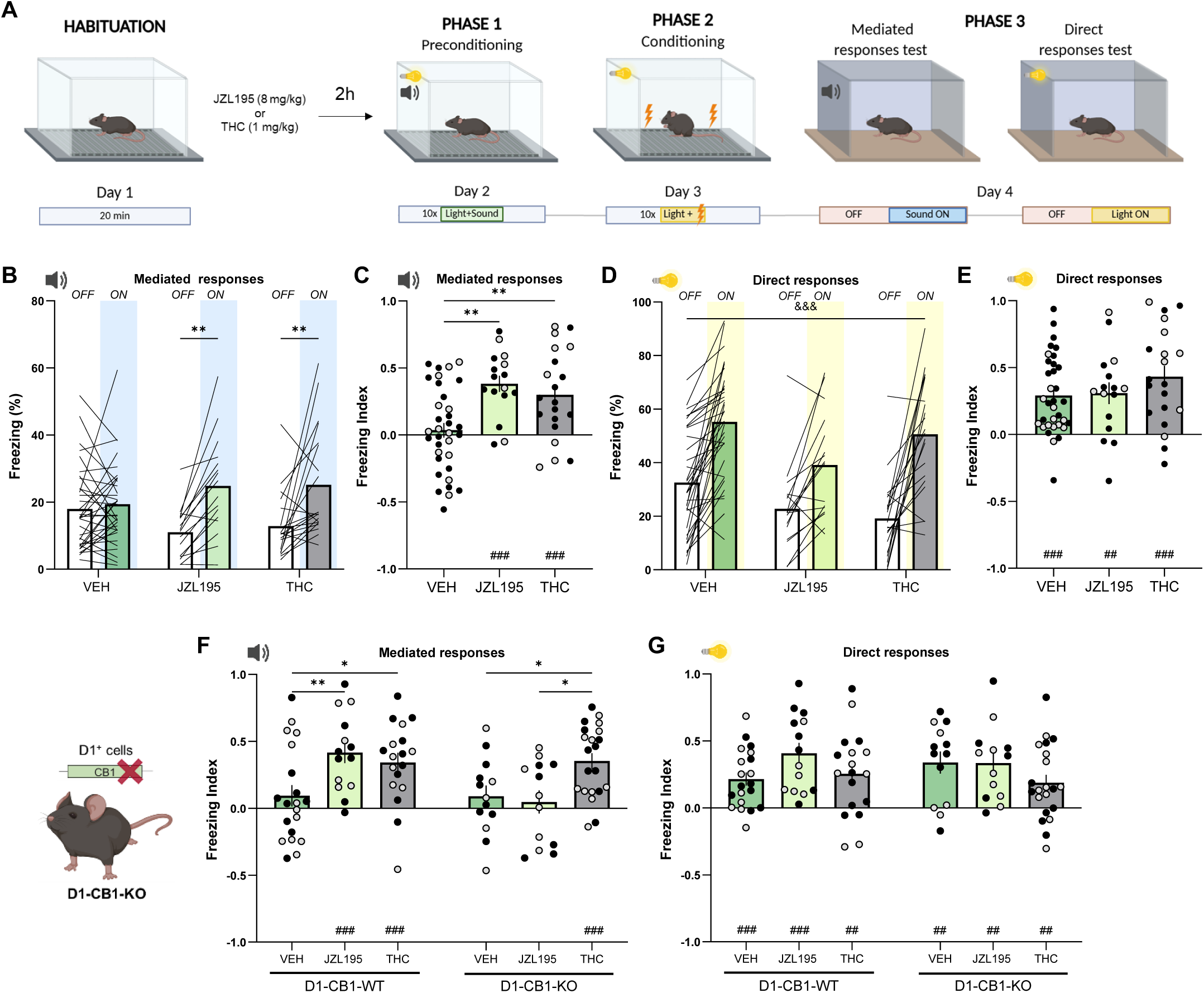
Endogenous and exogenous cannabinoids facilitate mediated responses through different mechanisms. (A) Schematic representation of the light and sound sensory preconditioning task with the JZL195 (8 mg/kg) or THC (1 mg/kg) injections prior to each preconditioning session. (B-G) (B) Freezing percentage of the JZL195, THC and vehicle groups in the last 60 seconds of the OFF period and the first 60 seconds of the ON period in the sound test. Repeated measures two-way ANOVA, interaction p=0,0027; Sidak’s post-hoc, p(ON vs OFF in the JZL195 group)=0,0058, p(ON vs OFF in the THC group)=0,0059. (C) Freezing index of the JZL195, THC and vehicle groups in the mediated responses test. Ordinary one-way ANOVA, p=0.0005; Tukey’s post-hoc, p(VEH vs JZL195)=0.0012, p(VEH vs THC)=0.0093. One sample t-test, p(JZL195)<0.0001, p(THC)=0.0008. (D) Freezing percentage of the JZL195, THC and vehicle groups in the last 60 seconds of the OFF period and the first 60 seconds of the ON period in the direct responses test. Repeated measures two-way ANOVA, ON vs OFF effect p<0.0001. (E) Freezing index of the JZL195, THC and vehicle groups in the direct responses test. One sample t-test, p(VEH)<0.0001, p(JZL195)=0.0017, p(THC)<0.0001. n_(Vehicle)_= 21M/13F, n_(JZL195)_= 11M/5F, n_(THC)_= 12M/8F. (F and G) Freezing index of the D1-CB1-WT and D1-CB1-KO mice after JZL195, THC and vehicle treatments during preconditioning in the mediated (F) and direct responses (G) tests. (F) Two-way ANOVA, interaction p=0.0307; Tukey’s post-hoc, p(D1-CB1-WT VEH vs JZL195)=0.0081, p(D1-CB1-WT VEH vs THC)=0.0375, p(D1-CB1-KO VEH vs THC)=0.0433, p(D1-CB1-KO JZL195 vs THC)=0.0154. One sample t-test, p(D1-CB1-WT, JZL)<0.0001, p(D1-CB1-WT, THC)=0.0002, p(D1-CB1-KO, THC)<0.0001. (G) One sample t-test, p(D1-CB1-WT, VEH)=0.0003, p(D1-CB1-WT, JZL)=0.0014, p(D1-CB1-WT, THC)=0.0002, p(D1-CB1-KO, VEH)=0.001, p(D1-CB1-KO, JZL)=0.0036, p(D1-CB1-KO, THC)=0.0058. n_(D1-CB1-WT,Vehicle)_= 9M/11F, n_(D1-CB1-WT,JZL195)_= 8M/6F, n_(D1-CB1-WT,THC)_= 10M/8F, n_(D1-CB1-KO,Vehicle)_= 8M/5F, n_(D1-CB1-KO,JZL195)_= 6M/7F, n_(D1-CB1-KO,THC)_= 10M/11F. Black dots: males; grey dots: females. Data are expressed in mean ± SEM. *p<0.05, **p<0.01, ***p<0.001, ^###^p<0.001, ^&&&^p<0.001.

To further elucidate if this potentiation involved CB1 receptors in D1 positive cells, we applied the 10 light-sound pairings protocol with JZL195 or THC to D1-CB1-KO mouse line. All groups showed direct responses to light (Fig. 4G and Supplementary Fig. 4E), but mediated responses yielded divergent results. Whereas THC-induced mediated responses were still present in D1-CB1-KO mice, mediated responses promoted by JZL195 (present in D1-CB1-WT mice) was abolished in D1-CB1-KO mice (Fig. 4F and Supplementary Fig. 4C). These results indicate that cannabinoid facilitation of IA memory formation during preconditioning occurs through distinct mechanisms depending on the origin of cannabinoid signaling. While JZL195 requires CB1 receptors in D1-positive cells for its effect, THC requires CB1 receptors other than those located in D1-positive cells. We therefore concluded that the physiological process underlying IA memory formation, both in the standard preconditioning protocol and in a shorter protocol with increased endocannabinoid tone, requires CB1 receptors in D1-positive cells.

## DISCUSSION

Our daily choices are shaped by prior stimulus-reinforcer associations, as well as by unreinforced or incidental associations (IAs) that occur between neutral stimuli (Johansen et al., 2011; Wimmer & Shohamy, 2012). Despite their relevance for adaptability and survival, IAs remain comparatively understudied, a gap that has motivated growing efforts to uncover their underlying brain mechanisms (Gewirtz & Davis, 2000; Parkes & Westbrook, 2011). Researchers have successfully used sensory preconditioning tasks that allow mediated learning and mediated responses to be investigated during conditioning and retrieval tests, respectively (Holmes et al., 2013, 2018; Sadacca et al., 2016; Taylor-Yeremeeva et al., 2021; Wong et al., 2019, 2026), yet few studies have addressed IA formation itself, occurring during preconditioning (Busquets-Garcia et al., 2018; Sadacca et al., 2018; Talaron et al., 2026). Using a recently developed light-sound preconditioning protocol in mice (Pinho et al., 2025), we obtained robust mediated responses in both male and female mice only when light and sound are initially paired, demonstrating the associative nature of the phenomenon instead of generalization (Holmes et al., 2022). Although the initial study reported sex differences in this paradigm, with reliable mediated responses observed only in males (Pinho et al., 2025), our data show comparable mediated responses in male and female mice. This discrepancy may stem from methodological differences, such as the inverted light/dark cycle or the intensity of the light and sound stimuli used, illustrating that IA formation is a very sensitive process that can be easily altered due to changes in experimental conditions.

Previous work showed that systemic administration of the CB1R receptor antagonist during S1-S2 pairings blocked subsequent mediated responses using either an aversive odor-taste preconditioning task or an appetitive light-sound preconditioning paradigm (Busquets-Garcia et al. 2018). Here we found that hippocampal CB1 receptors are also required for light-sound preconditioning as previously obtained with odor-taste preconditioning (Busquets-Garcia et al., 2018). Together, these findings, indicating that IA memory formation relies on a shared hippocampal mechanism regardless of sensory modalities (Busquets-Garcia et al., 2018; Iordanova et al., 2011; Lin et al., 2016; Port et al., 1987; Talaron et al., 2026; Voss et al., 2017; Wheeler et al., 2013; Wimmer & Shohamy, 2012).

Dopamine signaling in several brain regions different from the hippocampus has been proposed to play an important role in sensory preconditioning (Costa et al., 2025; Fry et al., 2020; Roughley et al., 2021; Sadacca et al., 2016; Seitz et al., 2021; Sharpe et al., 2017, 2020; Young et al., 1998). Notably, a recent study indicates that memory linking, a form of unreinforced memory that relies on contextual cues, depends on dopaminergic projections from the locus coeruleus to the hippocampus (Chowdhury et al., 2022). Here, we show that hippocampal dopamine signaling is required for IA formation. Our data reveal higher dopamine activity during light-sound pairings, relative to the unpaired condition, driven primarily by reduced hippocampal dopamine activity when sound was presented alone. This low hippocampal dopamine response to sound is consistent with previous work showing that auditory-evoked dopamine activity typically requires sound intensities of at least 85 dB (Wilmot et al., 2024), well above the 50 dB used here. This group difference did not extend to the conditioning phase, where value is assigned to light through footshock pairing. Importantly, during the mediated responses test, the paired group showed a higher dopamine peak upon sound presentation, resembling dopamine activity evoked by light. This indicates that light-sound pairings during preconditioning induced long-lasting hippocampal plasticity, enabling a stimulus that did not evoke a response beforehand to now enhance dopamine activity. Moreover, this differential dopamine activity between paired and unpaired groups was specific to the sound as light presentation elicited similar dopamine activity in both groups. We then investigated if dopamine type-1 receptors (D1) in the hippocampus would play a role in IA formation. We show that pharmacological blockade of these receptors prior to each preconditioning session was sufficient to block mediated responses in the sound test. Although we cannot directly attribute this blockade to hippocampal D1 receptors, the fact that hippocampal dopamine signaling is necessary for light-sound preconditioning suggests that systemic D1 blockade will likely disrupt hippocampal dopaminergic signaling.

Given that both CB1 receptors and dopaminergic signaling are independently required for light-sound preconditioning, we next asked whether these two systems converge onto a common neuronal population to support IA memory formation. Using a mouse line lacking CB1 receptors specifically in D1-positive cells (D1-CB1-KO; Monory et al., 2007), we found impaired mediated responses in both the light-sound and the odor-taste preconditioning protocols, demonstrating that CB1 receptors and dopaminergic signaling interact through CB1 receptors located in D1-positive cells to form IAs across sensory modalities. Whether this D1-CB1 population is specifically located within the hippocampus remains to be directly tested. However, the fact that hippocampal CB1 receptors are required for IAs regardless of sensory pairings, is consistent with this possibility. The population of hippocampal D1- and CB1-positive cells is broad, spanning from glutamatergic pyramidal cells to GABAergic interneurons (Puighermanal et al., 2017). Previous work on odor-taste preconditioning found that CB1 receptors located specifically on hippocampal GABAergic interneurons are required for IA (Busquets-Garcia et al., 2018), and a subpopulation of these interneurons have been identified to co-express D1 receptors (Oliveira Da Cruz et al., 2020; Puighermanal et al., 2017). Together, these data points to CB1 receptors expressed in hippocampal D1-positive interneurons to be key regulators of IAs.

Having established that CB1 receptors in D1-positive cells are necessary for IA memory formation under standard preconditioning conditions, we then wondered whether activation of CB1 receptor could promote mediated responses. Previous studies indicate that a minimal number of S1-S2 pairings during preconditioning is necessary to obtain subsequent mediated responses (Busquets-Garcia, Soria-Gómez, Redon, et al., 2017; Hoffeld et al., 1960), and CB1 receptor protein expression in the hippocampus is related to the number of odor-taste pairings (Busquets-Garcia et al., 2018). Using suboptimal preconditioning based on 10 light-sound pairings, instead of the usual 30 pairings, we pharmacologically enhance CB1 receptor activity thanks to use of JZL195, an inhibitor of endocannabinoid degrading enzymes, or of cannabis-derived THC. Both treatments prior to preconditioning were sufficient to induce subsequent mediated responses under conditions that were insufficient for vehicle-treated mice. These findings indicate that CB1 receptor activation is sufficient to promote mediated responses, reinforcing the idea that ECS is a key regulator of IAs. We then wondered whether this pharmacological facilitation could also be dependent on CB1 receptors located on D1-positive cells. Surprisingly, we found a dichotomous effect: whereas THC was still able to promote mediated responses in D1-CB1-KO mice, the promoting effect of JZL195 was abolished. This indicates that the facilitatory effect of JZL195 on IA memory formation depends on CB1 receptors located in D1-positive cells, whereas the effect of THC requires CB1 receptors expressed on other cell types. Taken together, these results indicate that physiological IA formation, under standard or suboptimal preconditioning conditions, requires CB1 receptors located in D1-positive cells.

In conclusion, our work demonstrates that hippocampal CB1 receptors are essential for IA memory formation across sensory modalities in both male and female mice. In addition, we show that dopaminergic signaling, which interacts with the ECS through a population of CB1- and D1-receptor-expressing cells, plays a critical role in this process. Finally, we demonstrate that endogenous and exogenous CB1 receptor activation can both promote IA formation under otherwise insufficient preconditioning conditions, acting through distinct mechanisms depending on whether cannabinoid tone is increased endogenously or exogenously. Together, these findings establish the ECS, and its interaction with hippocampal dopaminergic signaling, as a central mechanism governing IA memory formation across sensory modalities. This opens new venues to further explore the way cannabinoids control this process which might be key to unravel how disrupted IAs underlie some psychiatric conditions such as psychosis (Busquets-Garcia, Soria-Gómez, Ferreira, et al., 2017; Busquets-Garcia, Soria-Gómez, Redon, et al., 2017; Fry et al., 2020, 2024; M. A. McDannald et al., 2011; M. McDannald & Schoenbaum, 2009).

## ACKNOWLEDGEMENTS

We would like to thank the animal caretakers for mouse care and the platforms of the Neurocentre Magendie for genotyping of the samples. We thank the members of the Marsicano lab for enriching discussion and support. We would also like to thank Pierre Trifilieff and Lola Hardt for providing us the Dopamine sensor used in our fiber photometry experiments. This work was supported by French National Research Agency (ERA-Net Neuron CanShank, ANR-21-NEU2-0001-04, to G.M; CaMeLS, ANR-23-CE16-0022-01 to G.M.; Hippobese, ANR-23-CE14-0004-03 to G.F. and G.M.; F_r_IEND-24-CE17-XXX, to G.F. and G.M). U.B.F. was the recipient of a PhD fellowship (2023-2025) and an extension grant (2025-2026) from the Foundation pour la Recherche Medicale (FDT202504020246). E.R. was the recipient of a PhD fellowship (2025-2028) from the Bordeaux NeuroCampus Graduate Program, managed by the French National Research Agency (ANR-17-EURE-0028).

## AUTHOR CONTRIBUTIONS

G.M. and G.F contributed to the conception of the project. U.B.F., M.B.-C., E.R. and C.I. performed and analyzed the experiments. P.G.-S. developed the Python scripts used for behavioral and fiber photometry analysis. S.B. helped with fiber photometry experiments. J.P. and M.G.-P. helped setting up the light-sound preconditioning protocol. A.B.-G. helped setting up the odor-taste preconditioning protocol. U.B.F, M.B.-C, G.M. and G.F redacted the manuscript.

