## Supplementary Figures for "Endocannabinoid-dopamine interactions mediate incidental associations in the hippocampus"

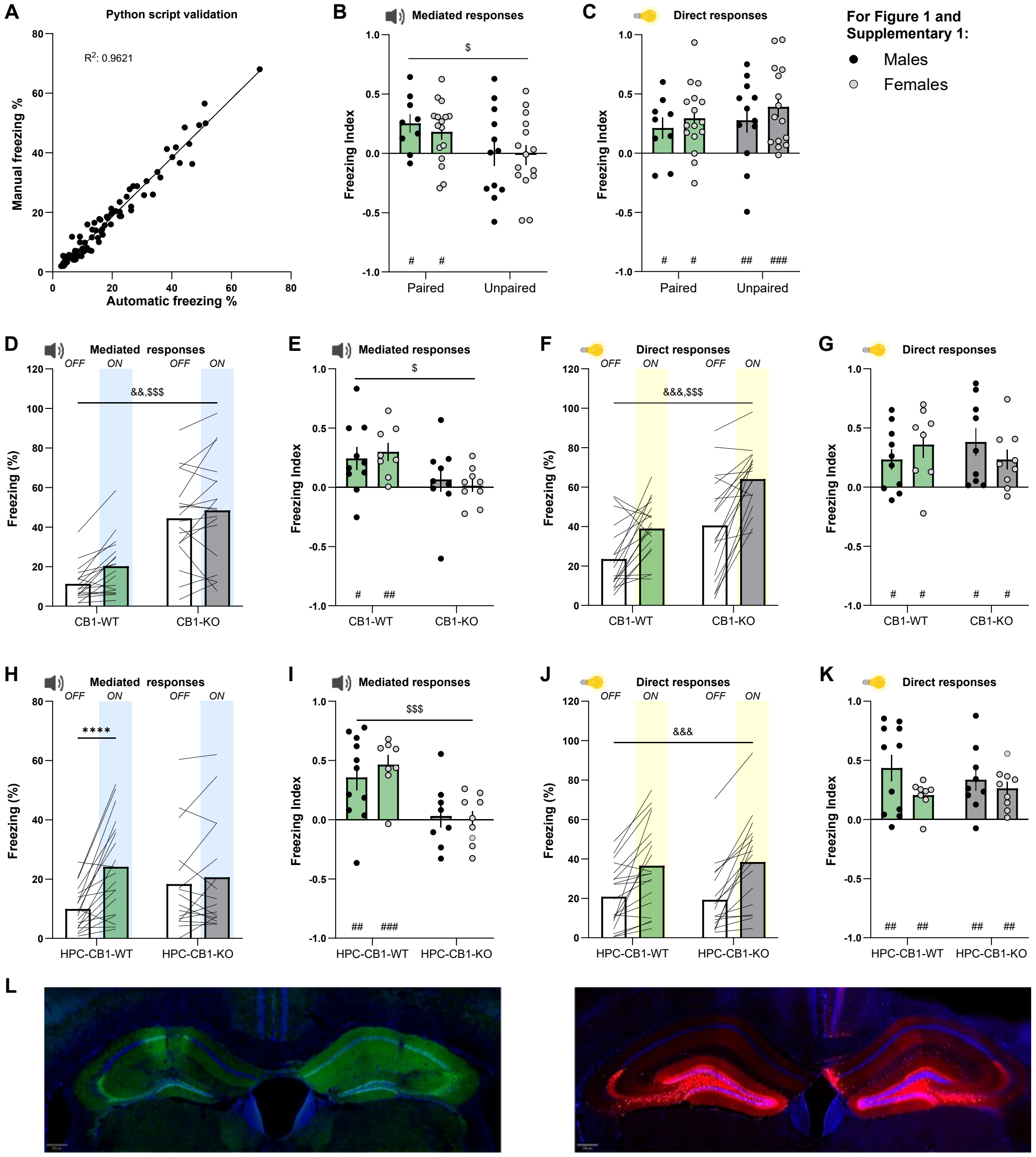


**Supplementary Figure 1: Hippocampal CB1 receptors are required for light sound sensory preconditioning.**

(A) Linear correlation between the manual and automatic freezing counting in the OFF and ON periods of the mediated and direct responses test of the male mice from Figure 1A-G.

(B and C) Freezing index of the male and female mice in the paired and unpaired groups in the mediated responses (B) and direct responses (C) tests. (B) Two-way ANOVA, paired vs unpaired effect p=0,0135. One sample t-test, p(paired males)=0.012, p(paired females)=0.0138. (C) One sample t-test, p(paired males)=0.044, p(paired females)=0.022, p(unpaired males)=0.001, p(paired females)=0.0005. $n_{(paired)}$= 9M/16F, $n_{(unpaired)}$= 12M/16F.

(D-G) (D) Freezing percentage of the CB1-WT and CB1-KO mice in the last 60 seconds of the OFF period and the first 60 seconds of the ON period in the sound test. Repeated measures two-way ANOVA, ON vs OFF effect p=0,0014, CB1-WT vs CB1-KO effect p<0.0001.

(E) Freezing index of the male and femaleCB1-WT and CB1-KO mice in the mediated responses test after paired preconditioning. Two-way ANOVA, CB1-WT vs CB1-KO effect p=0,0115. One sample t-test, p(CB1-WT males)=0.0321, p(CB1-WT females)=0.0056.

(F) Freezing percentage of the CB1-WT and CB1-KO mice in the last 60 seconds of the OFF period and the first 60 seconds of the ON period in the light test. Repeated measures two-way ANOVA, ON vs OFF effect p<0.0001, CB1-WT vs CB1-KO effect p=0.0005.

(G) Freezing index of the male and femaleCB1-WT and CB1-KO mice in the direct responses test after paired preconditioning. One sample t-test, p(CB1-WT males)=0.0237, p(CB1-WT females)=0.0146, p(CB1-KO males)=0.0122, p(CB1-KO females)=0.0244. $n_{(CB1\text{-}WT)}$= 10M/8F, $n_{(CB1\text{-}KO)}$= 9M/9F.

(H-K) (H) Freezing percentage of the HPC-CB1-WT and HPC-CB1-KO groups in the last 60 seconds of the OFF period and the first 60 seconds of the ON period in the sound test. Repeated measures two-way ANOVA, interaction p=0.003; Sidak's post-hoc, p(ON vs OFF in the HCP-CB1-WT group)<0.0001.

(I) Freezing index of the male and female HPC-CB1-WT and HPC-CB1-KO mice in the mediated responses test. Two-way ANOVA, HPC-CB1-WT vs HPC-CB1-KO effect p=0,0002. One sample t-test, p(HPC-CB1-WT males)=0.0081, p(HPC-CB1-WT females)=0.0007.

(J) Freezing percentage of the HPC-CB1-WT and HPC-CB1-KO groups in the last 60 seconds of the OFF period and the first 60 seconds of the ON period in the light test. Repeated measures two-way ANOVA, ON vs OFF effect p<0.0001.

(K) Freezing index of the male and female HPC-CB1-WT and HPC-CB1-KO mice in the direct responses test after paired preconditioning. One sample t-test, p(HPC-CB1-KO males)=0.0068, p(HPC-CB1-KO females)=0.0017. One sample Wilcoxon test p(HPC-CB1-WT males)=0.0028, p(HPC-CB1-WT females)=0.0017. $n_{(HPC\text{-}CB1\text{-}WT)}$= 11M/8F, $n_{(HPC\text{-}CB1\text{-}KO)}$= 8M/9F.

(L) Representative images showing hippocampal expression AAV-hSyn-CRE-GFP virus in HPC-CB1-KO (left) and AAV-hSyn-mCherry in HPC-CB1-WT mice. Scale bar, 250 µm.

Black dots: males; grey dots: females. Data are expressed in mean ± SEM. ****p<0.001, $^{\#}$p<0.05, $^{\#\#}$p<0.01, $^{\#\#\#}$p<0.001, $^{\&\&}$p<0.01, $^{\&\&\&}$p<0.001, $^{\$}$p<0.05, $^{\$\$\$}$p<0.001.

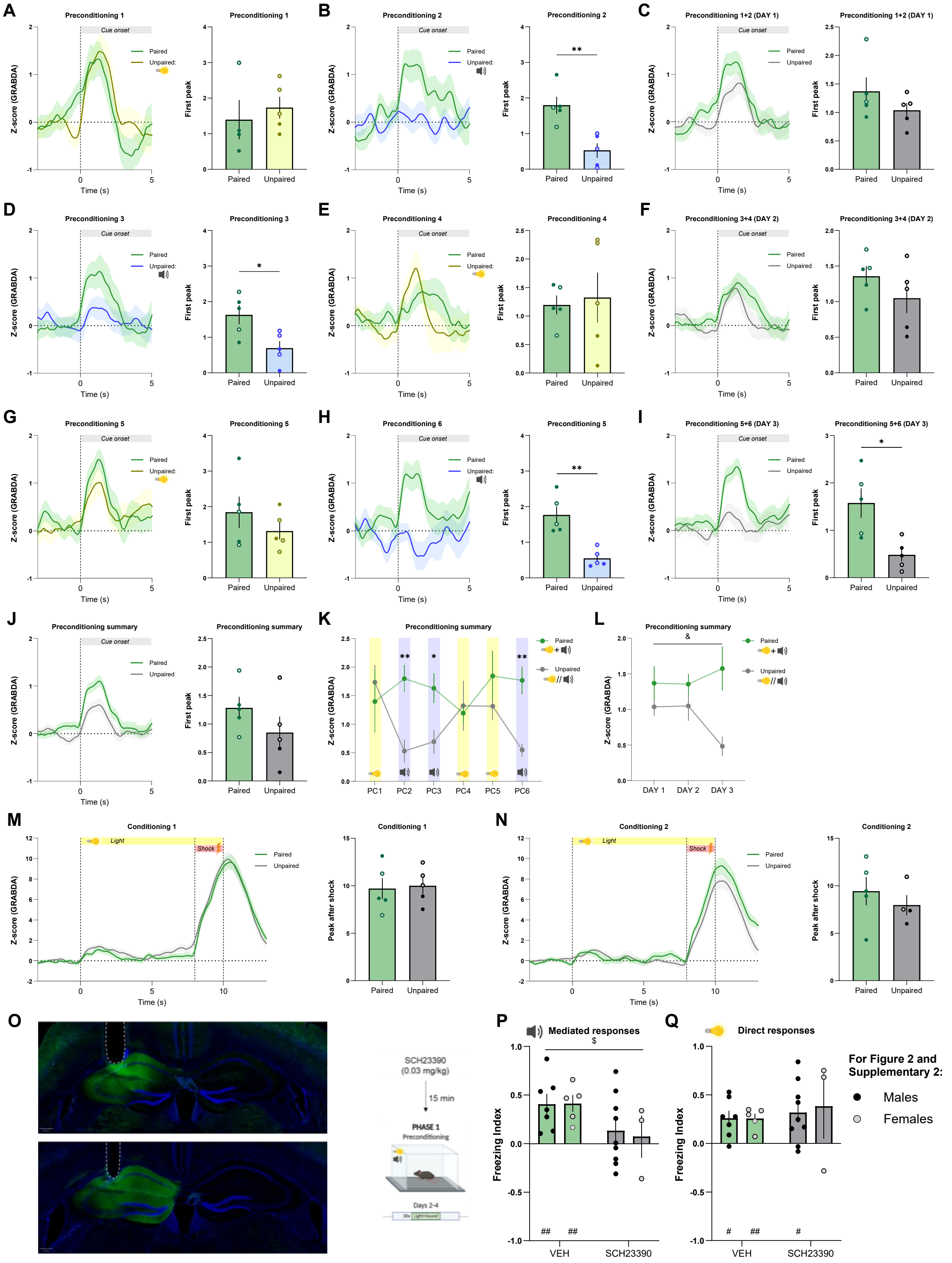
A
Preconditioning 1
Cue onset
Paired
Unpaired:
Z-score (GRABDA)
Time (s)
First peak
B
Preconditioning 2
C
Preconditioning 1+2 (DAY 1)
D
Preconditioning 3
E
Preconditioning 4
F
Preconditioning 3+4 (DAY 2)
G
Preconditioning 5
H
Preconditioning 6
I
Preconditioning 5+6 (DAY 3)
J
Preconditioning summary
K
PC1
PC2
PC3
PC4
PC5
PC6
L
DAY 1
DAY 2
DAY 3
M
Conditioning 1
Light
Shock
Peak after shock
N
Conditioning 2
O
SCH23390
(0.03 mg/kg)
15 min
PHASE 1
Preconditioning
Days 2-4
P
Mediated responses
Freezing Index
VEH
SCH23390
Q
Direct responses
For Figure 2 and
Supplementary 2:
Males
Females

**Supplementary Figure 2: Hippocampal dopamine signaling is necessary to form IAs.**

(A-N) (A) Average Z-scored ΔF/F Dopamine responses in neurons of the hippocampus aligned to the 5 light and sound simultaneous presentation onset in the preconditioning session 1 in mice following paired preconditioning, or aligned to the 5 light presentations in mice following unpaired preconditioning (left). First peak of average Z-scored ΔF/F Dopamine responses in the paired and unpaired (light) groups (right).

(B) Average Z-scored ΔF/F Dopamine responses in neurons of the hippocampus aligned to the 5 light and sound simultaneous presentation onset in the preconditioning session 2 in mice following paired preconditioning, or aligned to the 5 sound presentations in mice following unpaired preconditioning (left). First peak of average Z-scored ΔF/F Dopamine responses in the paired and unpaired (sound) groups (right). Unpaired t-test, $p=0.0035$.

(C) Average Z-scored ΔF/F Dopamine responses in neurons of the hippocampus aligned to the 10 light and sound simultaneous presentation onset in preconditioning day 1 of mice following paired preconditioning, or aligned to the 5 light and 5 sound presentations in mice following unpaired preconditioning (left). First peak of average Z-scored ΔF/F Dopamine responses in the paired and unpaired groups (right).

(D) Average Z-scored ΔF/F Dopamine responses in neurons of the hippocampus aligned to the 5 light and sound simultaneous presentation onset in the preconditioning session 3 in mice following paired preconditioning, or aligned to the 5 sound presentations in mice following unpaired preconditioning (left). First peak of average Z-scored ΔF/F Dopamine responses in the paired and unpaired (sound) groups (right). Unpaired t-test, $p=0.0213$.

(E) Average Z-scored ΔF/F Dopamine responses in neurons of the hippocampus aligned to the 5 light and sound simultaneous presentation onset in the preconditioning session 4 in mice following paired preconditioning, or aligned to the 5 light presentations in mice following unpaired preconditioning (left). First peak of average Z-scored ΔF/F Dopamine responses in the paired and unpaired (light) groups (right).

(F) Average Z-scored ΔF/F Dopamine responses in neurons of the hippocampus aligned to the 10 light and sound simultaneous presentation onset in preconditioning day 2 of mice following paired preconditioning, or aligned to the 5 light and 5 sound presentations in mice following unpaired preconditioning (left). First peak of average Z-scored ΔF/F Dopamine responses in the paired and unpaired groups (right).

(G) Average Z-scored ΔF/F Dopamine responses in neurons of the hippocampus aligned to the 5 light and sound simultaneous presentation onset in the preconditioning session 5 in mice following paired preconditioning, or aligned to the 5 light presentations in mice following unpaired preconditioning (left). First peak of average Z-scored ΔF/F Dopamine responses in the paired and unpaired (light) groups (right).

(H) Average Z-scored ΔF/F Dopamine responses in neurons of the hippocampus aligned to the 5 light and sound simultaneous presentation onset in the preconditioning session 6 in mice following paired preconditioning, or aligned to the 5 sound presentations in mice following unpaired preconditioning (left). First peak of average Z-scored ΔF/F Dopamine responses in the paired and unpaired (sound) groups (right). Unpaired t-test, $p=0.0017$.

(I) Average Z-scored ΔF/F Dopamine responses in neurons of the hippocampus aligned to the 10 light and sound simultaneous presentation onset in preconditioning day 3 of mice following paired preconditioning, or aligned to the 5 light and 5 sound presentations in mice following unpaired preconditioning (left). First peak of average Z-scored ΔF/F Dopamine responses in the paired and unpaired groups (right). Unpaired t-test, p=0.0121.

(J) Average Z-scored ΔF/F Dopamine responses in neurons of the hippocampus aligned to the 30 light and sound simultaneous presentation onset in all the preconditioning sessions of mice following paired preconditioning, or aligned to the 15 light and 15 sound presentations in mice following unpaired preconditioning (left). First peak of average Z-scored ΔF/F Dopamine responses in the paired and unpaired groups (right).

(K) Timeline of the first peaks of average Z-scored ΔF/F Dopamine responses along the preconditioning sessions. Two-way ANOVA, interaction p=0.02; Tukey's post-hoc p(paired vs unpaired in PC2)=0.0037, p(paired vs unpaired in PC3)=0.0291, p(paired vs unpaired in PC6)=0.0051.

(L) Timeline of the first peaks of average Z-scored ΔF/F Dopamine responses along the preconditioning days. Two-way ANOVA, paired vs unpaired effect p=0.0143.

(M) Average Z-scored ΔF/F Dopamine responses aligned to the 5 light and shock presentation onset during conditioning session 1 (left). First peak of average Z-scored ΔF/F Dopamine responses in the paired and unpaired groups after the footshock in conditioning session 1 (right).

(N) Average Z-scored ΔF/F Dopamine responses aligned to the 5 light and shock presentation onset during conditioning session 2 (left). First peak of average Z-scored ΔF/F Dopamine responses in the paired and unpaired groups after the footshock in conditioning session 2 (right). $n_{(paired)}$= 3M/2F, $n_{(unpaired)}$= 2M/3F.

(O) Representative images showing GRAB(DA2m) sensor endogenous fluorescence and optic fiber placement in mice following paired (top) and unpaired (bottom) preconditioning. Scale bar, 250 µm.

(P and Q) Freezing index of male and female SCH23390 (0.03 mg/kg) and vehicle-treated mice in the mediated responses (P) and direct responses (Q) tests after paired preconditioning. (P) Two-way ANOVA, vehicle vs SCH23390 effect p=0,0381. One sample t-test, p(vehicle males)=0.0079, p(vehicle females)=0.0088. (Q) One sample t-test, p(vehicle males)=0.0141, p(vehicle females)=0.0074, p(SCH23390 males)=0.0152. $n_{(vehicle)}$= 7M/5F, $n_{(SCH23390)}$= 9M/3F.

Black and filled dots: males; grey dots: females. Data are expressed in mean ± SEM. *p<0.05, **p<0.01, $^{\#}$p<0.05, $^{\#\#}$p<0.01, $^{\&}$p<0.05, $^{\$}$p<0.05.

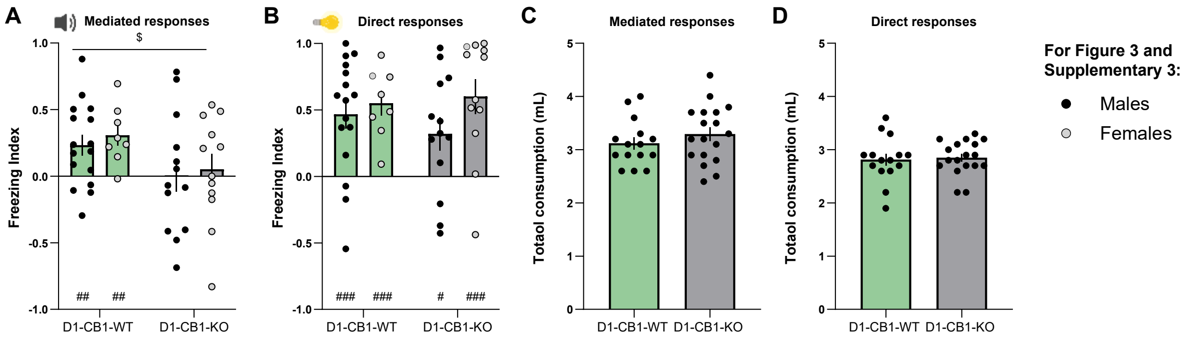


**Supplementary Figure 3: CB1 receptors in D1-positive cells are required for sensory preconditioning.**

(A and B) Freezing index of male and female D1-CB1-WT and D1-CB1-KO mice in the mediated responses (A) and direct responses (B) tests after light and sound preconditioning. (A) Two-way ANOVA, D1-CB1-WT vs D1-CB1-KO effect p=0,0312. One sample t-test, p(D1-CB1-WT males)=0.0091, p(D1-CB1-WT females)=0.0055. (B) One sample t-test, p(D1-CB1-WT males)=0.0007, p(D1-CB1-WT females)<0.0001, p(D1-CB1-KO males)=0.0254, p(D1-CB1-KO females)=0.0003. $n_{(D1\text{-}CB1\text{-}WT)}$= 16M/8F, $n_{(D1\text{-}CB1\text{-}KO)}$= 13M/12F.

(C and D) Total liquid consumption of D1-CB1-WT and D1-CB1-KO mice in the taste test (C) and odor test (D). $n_{(D1\text{-}CB1\text{-}WT)}$= 15M, $n_{(D1\text{-}CB1\text{-}KO)}$= 18M.

Black dots: males; grey dots: females. Data are expressed in mean ± SEM. $^{\#}p<0.05$, $^{\#\#}p<0.01$, $^{\#\#\#}p<0.001$, $^{\$}p<0.05$.

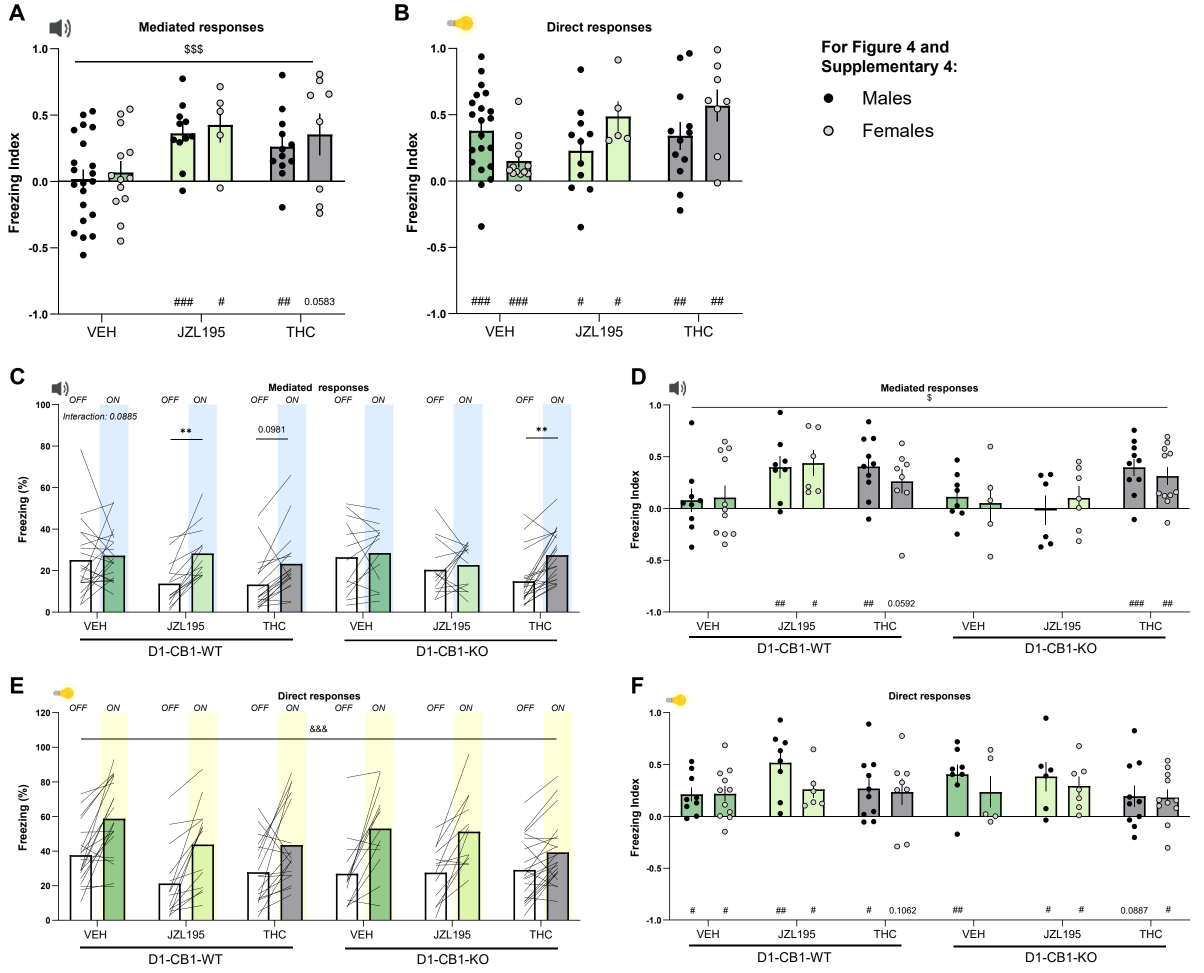


**Supplementary Figure 4: Endogenous and exogenous cannabinoids facilitate mediated responses through different mechanisms.** (A and (A and B) Freezing index of male and female the JZL195, THC and vehicle-treated mice in the mediated responses (A) and direct responses (B) tests. (A) Two-way ANOVA, treatment effect p=0.0009. One sample t-test, p(vehicle males)=0.0079, p(vehicle females)=0.0088. (B) One sample t-test, p(JZL195 males)=0.0004, p(JZL195 females)=0.0325, p(THC males)=0.0044, p(THC females)=0.0583. $n_{(Vehicle)}$= 21M/13F, $n_{(JZL195)}$= 11M/5F, $n_{(THC)}$= 12M/8F.

(C-F) (C) Freezing percentage of the D1-CB1-WT and D1-CB1-KO mice under JZL195, THC and vehicle treatments in the last 60 seconds of the OFF period and the first 60 seconds of the ON period in the sound test. Repeated measures three-way ANOVA, interaction p(Treatment x Genotype x ON vs OFF)=0,0885; Tukey's post-hoc, p(ON vs OFF in the D1-CB1-WT, JZL195 group)=0,0072, p(ON vs OFF in the D1-CB1-WT, THC group)=0,0981, p(ON vs OFF in the D1-CB1-KO, THC group)=0,0033.

(D) Freezing index of male and female D1-CB1-WT and D1-CB1-KO mice under JZL195, THC and vehicle treatments in the mediated responses test. Three-way ANOVA, Treatment x Genotype effect p=0,0308. One sample t-test, p(D1-CB1-WT, JZL195, males)=0.0075, p(D1-CB1-WT,

JZL195, females)=0.0165, p(D1-CB1-WT, THC, males)=0.0016, , p(D1-CB1-WT, THC, females)=0.0592, , p(D1-CB1-KO, THC, males)=0.0009, p(D1-CB1-KO, THC, females)=0.0042.

(E) Freezing percentage of the D1-CB1-WT and D1-CB1-KO mice under JZL195, THC and vehicle treatments in the last 60 seconds of the OFF period and the first 60 seconds of the ON period in the light test. Repeated measures three-way ANOVA, ON vs OFF effect p<0.0001.

(F) Freezing index of male and female D1-CB1-WT and D1-CB1-KO mice under JZL195, THC and vehicle treatments in the direct responses test. One sample t-test, p(D1-CB1-WT, vehicle, males)=0.0119, p(D1-CB1-WT, vehicle, females)=0.0145, p(D1-CB1-WT, JZL195, males)=0.0021, p(D1-CB1-WT, JZL195, females)=0.0263, p(D1-CB1-WT, THC, males)=0.0208, p(D1-CB1-WT, THC, females)=0.1062, p(D1-CB1-KO, vehicle, males)=0.0038, p(D1-CB1-KO, JZL195, males)=0.0424, p(D1-CB1-KO, JZL195, males)=0.0151, p(D1-CB1-KO, THC, males)=0.0887, p(D1-CB1-KO, THC, females)=0.036. $n_{(D1\text{-}CB1\text{-}WT,Vehicle)}$= 9M/11F, $n_{(D1\text{-}CB1\text{-}WT,JZL195)}$= 8M/6F, $n_{(D1\text{-}CB1\text{-}WT,THC)}$= 10M/8F, $n_{(D1\text{-}CB1\text{-}KO,Vehicle)}$= 8M/5F, $n_{(D1\text{-}CB1\text{-}KO,JZL195)}$= 6M/7F, $n_{(D1\text{-}CB1\text{-}KO,THC)}$= 10M/11F.

Black dots: males; grey dots: females. Data are expressed in mean ± SEM. **p<0.01, $^{\#}$p<0.05, $^{\#\#}$p<0.01, $^{\#\#\#}$p<0.001, $^{\&\&\&}$p<0.001, $^{\$}$p<0.05, $^{\$\$\$}$p<0.001.

(E) Freezing percentage of the D1-CB1-WT and D1-CB1-KO mice under JZL195, THC and vehicle treatments in the last 60 seconds of the OFF period and the first 60 seconds of the ON period in the light test. Repeated measures three-way ANOVA, ON vs OFF effect p<0.0001.

(F) Freezing index of male and female D1-CB1-WT and D1-CB1-KO mice under JZL195, THC and vehicle treatments in the direct responses test. One sample t-test, p(D1-CB1-WT, vehicle, males)=0.0119, p(D1-CB1-WT, vehicle, females)=0.0145, p(D1-CB1-WT, JZL195, males)=0.0021, p(D1-CB1-WT, JZL195, females)=0.0263, p(D1-CB1-WT, THC, males)=0.0208, p(D1-CB1-WT, THC, females)=0.1062, p(D1-CB1-KO, vehicle, males)=0.0038, p(D1-CB1-KO, JZL195, males)=0.0424, p(D1-CB1-KO, JZL195, males)=0.0151, p(D1-CB1-KO, THC, males)=0.0887, p(D1-CB1-KO, THC, females)=0.036. $n_{(D1\text{-}CB1\text{-}WT,Vehicle)}$= 9M/11F, $n_{(D1\text{-}CB1\text{-}WT,JZL195)}$= 8M/6F, $n_{(D1\text{-}CB1\text{-}WT,THC)}$= 10M/8F, $n_{(D1\text{-}CB1\text{-}KO,Vehicle)}$= 8M/5F, $n_{(D1\text{-}CB1\text{-}KO,JZL195)}$= 6M/7F, $n_{(D1\text{-}CB1\text{-}KO,THC)}$= 10M/11F.

Black dots: males; grey dots: females. Data are expressed in mean ± SEM. **p<0.01, $^{\#}$p<0.05, $^{\#\#}$p<0.01, $^{\#\#\#}$p<0.001, $^{\&\&\&}$p<0.001, $^{\$}$p<0.05, $^{\$\$\$}$p<0.001.
